# Estrogen-induced chromatin looping changes identify a subset of functional regulatory elements

**DOI:** 10.1101/2024.06.12.598690

**Authors:** Hosiana Abewe, Alexandra Richey, Jeffery M Vahrenkamp, Matthew Ginley-Hidinger, Craig M Rush, Noel Kitchen, Xiaoyang Zhang, Jason Gertz

## Abstract

Transcriptional enhancers can regulate individual or multiple genes through long-range three-dimensional (3D) genome interactions, and these interactions are commonly altered in cancer. Yet, the functional relationship between changes in 3D genome interactions associated with regulatory regions and differential gene expression appears context-dependent. In this study, we used HiChIP to capture changes in 3D genome interactions between active regulatory regions of endometrial cancer cells in response to estrogen treatment and uncovered significant differential long-range interactions strongly enriched for estrogen receptor *α*(ER) bound sites (ERBS). The ERBS anchoring differential chromatin loops with either a gene’s promoter or distal regions were correlated with larger transcriptional responses to estrogen compared to ERBS not involved in differential 3D genome interactions. To functionally test this observation, CRISPR-based Enhancer-i was used to deactivate specific ERBS, which revealed a wide range of effects on the transcriptional response to estrogen. However, these effects are only subtly and not significantly stronger for ERBS in differential chromatin loops. In addition, we observed an enrichment of 3D genome interactions between the promoters of estrogen upregulated genes and found that looped promoters can work together cooperatively. Overall, our work reveals that estrogen treatment causes large changes in 3D genome structure in endometrial cancer cells; however, these changes are not required for a regulatory region to contribute to an estrogen transcriptional response.

## Introduction

Enhancers are critical *cis*-regulatory elements in metazoan genomes that increase the transcription of target genes in response to intrinsic and external signals. A typical human gene is associated with multiple enhancers, each bound by specific transcription factors (Furlong and Levine 2018; Andersson et al. 2014). These enhancers can reside far from their target gene promoters and interact through long-range chromatin looping events as part of the three-dimensional (3D) genome structure (Heidari et al. 2014). Aberrant 3D genome interactions, particularly involving Promoter–enhancer contacts, can lead to dysregulation of oncogenes and tumor suppressors (Valton and Dekker 2016). In fact, most changes in 3D genome structure have been reported in relatively small-scale chromatin loops, which have a size range of 5 -100 kb and primarily involve Promoter–enhancer and Enhancer–enhancer interactions (Yang et al. 2019; Taberlay et al. 2016; Flavahan et al. 2016; Braun et al. 2019; O’Mara et al. 2019) . Furthermore, these 3D genome alterations are seen across different types of cancer (Kantidze et al. 2019; Wu et al. 2017; Hnisz et al. 2016; Barutcu et al. 2015) with cancer and normal cells exhibiting different 3D genome structures (Achinger-Kawecka et al. 2016; Kloetgen et al. 2020; Li et al. 2021).

While chromatin looping is a common feature of active enhancers, changes in the 3D genome structure aren’t always associated with differential gene expression. Some studies have detected significant effects on gene regulation that accompany large changes to 3D genome organization (Kim et al. 2024; Smits et al. 2023; Franke et al. 2016; Lupiáñez et al. 2015; Pinoli et al. 2020) . In contrast, other studies have demonstrated that broad rewiring of 3D genome interactions, due to acute depletion of topologically associating domain (TAD) boundaries or TAD architectural proteins (e.g., CTCF or cohesin), were unexpectedly accompanied by modest effects on gene expression (Despang et al. 2019; Schwarzer et al. 2017; Nora et al. 2017; Rao et al. 2017). These seemingly contradictory results suggest that changes to the 3D genome structure impact gene expression to varying degrees depending on context. This inconsistency is also seen in cancer cells, where abnormal spatial chromatin contacts in association with an active oncogene do not necessarily correlate with changes in gene expression (Li et al. 2020). To better understand the functional relationship between chromatin looping and gene expression in response to oncogene activation in cancer cells, we investigated changes in the 3D genome structure and the transcriptional response to estrogen receptor *α* (ER) activation in endometrial cancer cells.

ER is an oncogenic transcription factor in breast and endometrial cancer that is activated by binding to estrogens, such as endogenous 17*β*-estradiol (E2) (Rodriguez et al. 2019; Shen et al. 2016; Zwart et al. 2011). ER binds predominantly to distal enhancers and primarily asserts its effects on transcription from a distance through chromatin interactions (Fullwood et al. 2009). Previous studies have revealed that activation of ER upon estrogen exposure is associated with global reorganization of 3D genome interactions in breast cancer cell lines (Mourad et al. 2014). Additionally, a strong correlation between ER and changes in chromatin looping interactions has been reported in both breast (Fullwood et al. 2009; Gomez Acuna et al. 2024) and endometrial (La Greca et al. 2022) cancer cells, suggesting that ER may play a role in mediating 3D genome structure alterations in response to estrogen treatment or that it may utilize changes in chromatin looping to impact transcription. However, it remains unclear whether 3D genome changes relate to the functional importance of ER-bound enhancers in terms of the transcriptional response to estrogen exposure. Moreover, the studies into estrogen and 3D genome structure changes have been focused mainly on breast cancer, while its effect in endometrial cancer, where we have previously shown that ER regulates different genes than in breast cancer (Rodriguez et al. 2020), is poorly understood.

Here, we dissected the functional role of 3D genome structure changes in gene regulation upon estrogen treatment in endometrial cancer cells. First, we identified changes in the 3D genome structure in response to E2 treatment by performing HiChIP, a high-resolution technique for analyzing 3D genome structure (Mumbach et al. 2017). We characterized how ER binding events relate to 3D genome structure changes and how these changes correspond to transcriptional responses. To determine the functional importance of specific regulatory regions, we used CRISPR-based Enhancer-i (Carleton et al. 2017) to block ER-bound enhancers from different classes of chromatin interactions or promoters that are part of an E2 upregulated gene cluster. With these studies, our main goal was to uncover how changes in the 3D genome structure relate to the functional importance of regulatory regions involved in the estrogen transcriptional response of endometrial cancer cells.

## Results

### The 3D genome structure of endometrial cancer cells is altered in response to estrogen exposure

To investigate the effects of ER activation on the 3D genome structure of endometrial cancer cells, we performed HiChIP in the context of 1-hour treatment with E2 or vehicle. We used an antibody against acetylation of lysine 27 of histone H3 (H3K27ac). Histone acetylation is important for E2-induced transcription regulation (Ruh et al. 1999), and focusing on H3K27ac allowed us to concentrate signal on active promoters and enhancers. These experiments were performed in two different ER expressing endometrial cancer cell lines, Ishikawa and HCI-EC-23 (Rush et al. 2022). The metrics for all libraries indicated high-quality data (Figures S1A,B). We identified 417,556 and 844,121 total 3D genome interactions in Ishikawa and HCI-EC-23 cell lines, respectively. Most of these interactions represented chromatin loops less than 100 kb apart in the linear genome with a median distance between anchors of 24.0 kb for Ishikawa and 33.6 kb for HCI-EC-23 (Figure 1A); the median anchor size was 2922 bp and 2801 bp for Ishikawa and HCI-EC-23, respectively.

**Figure 1.**
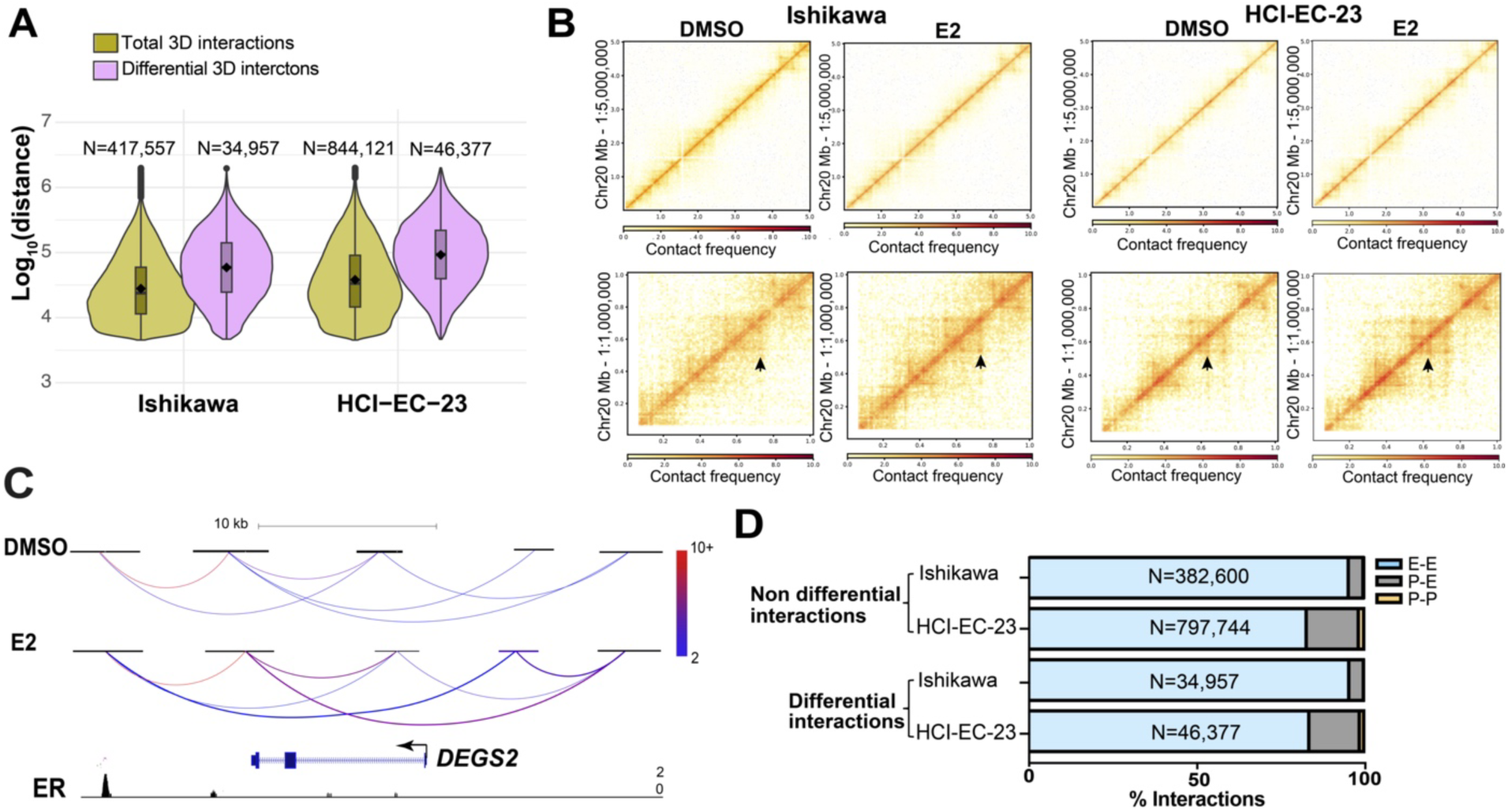
3D genome structure changes upon estrogen induction. A) Violin plots show distance between anchors for all 3D genome interactions (green) and differential interactions (purple) in Ishikawa (left) and HCI-EC-23 (right) cell lines identified by HiChIP. B) Heatmaps show portions of Chromosome 20 HiChIP signal for DMSO and E2 samples in Ishikawa (left) and HCI-EC-23 (right) cell lines (top panel represents lower resolution; bottom panel displays higher resolution). C) HiChIP 3D genome interactions at the *DEGS2* locus in Ishikawa cells are shown where the color of the chromatin loops represents the normalized read depth, and the two differential chromatin loops are shown as thicker in the E2 track. ChIP-seq browser track of ER upon E2 treatment is displayed at the bottom. D) Genomic distribution of non-differential and differential 3D genome interactions in Ishikawa and HCI-EC-23 cell lines are shown in terms of Promoter–promoter (P-P), Promoter–enhancer (P-E), and Enhancer–enhancer (E-E) interactions.

Differential analysis of HiChIP data identified 34,956 and 46,378 3D genome interactions impacted by E2 treatment (FDR < 0.05) in Ishikawa and HCI-EC-23 cell lines, respectively. Consistent with previous findings of E2 treatment in breast cancer cells (Zhou et al. 2019), the majority of identified differential interactions were longer than non-differential chromatin loops, representing long-range chromatin looping events (median size: 59.0 kb in Ishikawa and 94.0 kb in HCI-EC-23; Figure 1A), implying substantial involvement of distal enhancers in altering the 3D genome structure upon estrogen treatment. Taking Chromosome 20 as an example, the 3D genome structure did not change at low resolution but changed significantly at high resolution upon estrogen treatment (Figure 1B). Most of these observed changes involved increased interactions with E2 treatment rather than reduced interactions (68.8% and 69.9% of differential chromatin loops increased in Ishikawa and HCI-EC-23, respectively). As an example, *DEGS2* is an E2 upregulated gene that exhibited an increase in interaction frequencies with distal enhancers bound by ER upon estrogen treatment (Figure 1C). Interactions involving two distal enhancer regions predominated over Promoter–enhancer or Promoter–promoter interactions (Figure 1D), implying the involvement of distal enhancers in altering the 3D genome structure upon E2 induction. We have previously identified significant changes in H3K27ac after 8 hours of E2 treatment (Carleton et al. 2017); however, H3K27ac signal at peaks defined by self-ligation products was only impacted by the 1-hour E2 treatment at 1 region in Ishikawa cells and 2 regions in HCI-EC-23 cells, indicating that changes in H3K27ac are not responsible for the observed differential chromatin loops. Overall, this data provided a foundation for studying the relationship between chromatin looping and transcriptional responses to estrogen in endometrial cancer cells.

### Differential 3D genome interactions are associated with the estrogen transcriptional response

To understand how changes in 3D genome interactions in response to E2 treatment relate to gene expression, we first annotated differential 3D genome interactions that have anchors within 5 kb of transcription start sites (TSS). We identified 6027 and 11,023 genes that harbor differential interactions in response to E2 induction in Ishikawa and HCI-EC-23 cell lines, respectively. However, as the majority of differential interactions involve two distal enhancer regions, we also annotated differential chromatin interactions that have anchors within 100 kb of a TSS and found 18,672 and 19,746 genes associated with differential 3D genome interactions in response to E2 induction in Ishikawa and HCI-EC-23 cell lines, respectively. There was a significant overlap of genes with differential 3D genome interactions anchored within 5 kb (47.1%; *P*-value = 2.13x10^-69^, hypergeometric distribution) or 100 kb (92.3%; *P*-value = 7.86x10^-488^, hypergeometric distribution) of TSS between the two cell lines, suggesting that the endometrial cancer models consistently respond to E2 treatment in terms of 3D genome organization (Figures S1C, D). Estrogen induction has been previously shown to impact the expression of hundreds of genes in both endometrial cancer cell lines (Gertz et al. 2013; Rush et al. 2022), particularly in Ishikawa. Integration of E2 responsive genes with genes that have differential 3D genome interactions anchored within 5 kb of their TSS revealed that Ishikawa cells exhibited a significant enrichment of E2 regulated genes, particularly upregulated genes, in those involved with differential 3D genome interactions (Figure 2A; *P*-value = 0.001 for upregulated genes and 0.03 for downregulated genes, hypergeometric distribution). In contrast, HCI-EC-23 cells showed a weaker association between E2 regulated genes and differential 3D genome interactions (Figure S2A; *P*-value = 0.46 for upregulated genes and 0.30 for downregulated genes, hypergeometric distribution), which might be due to a lower number of E2 responsive genes in HCI-EC-23.

**Figure 2.**
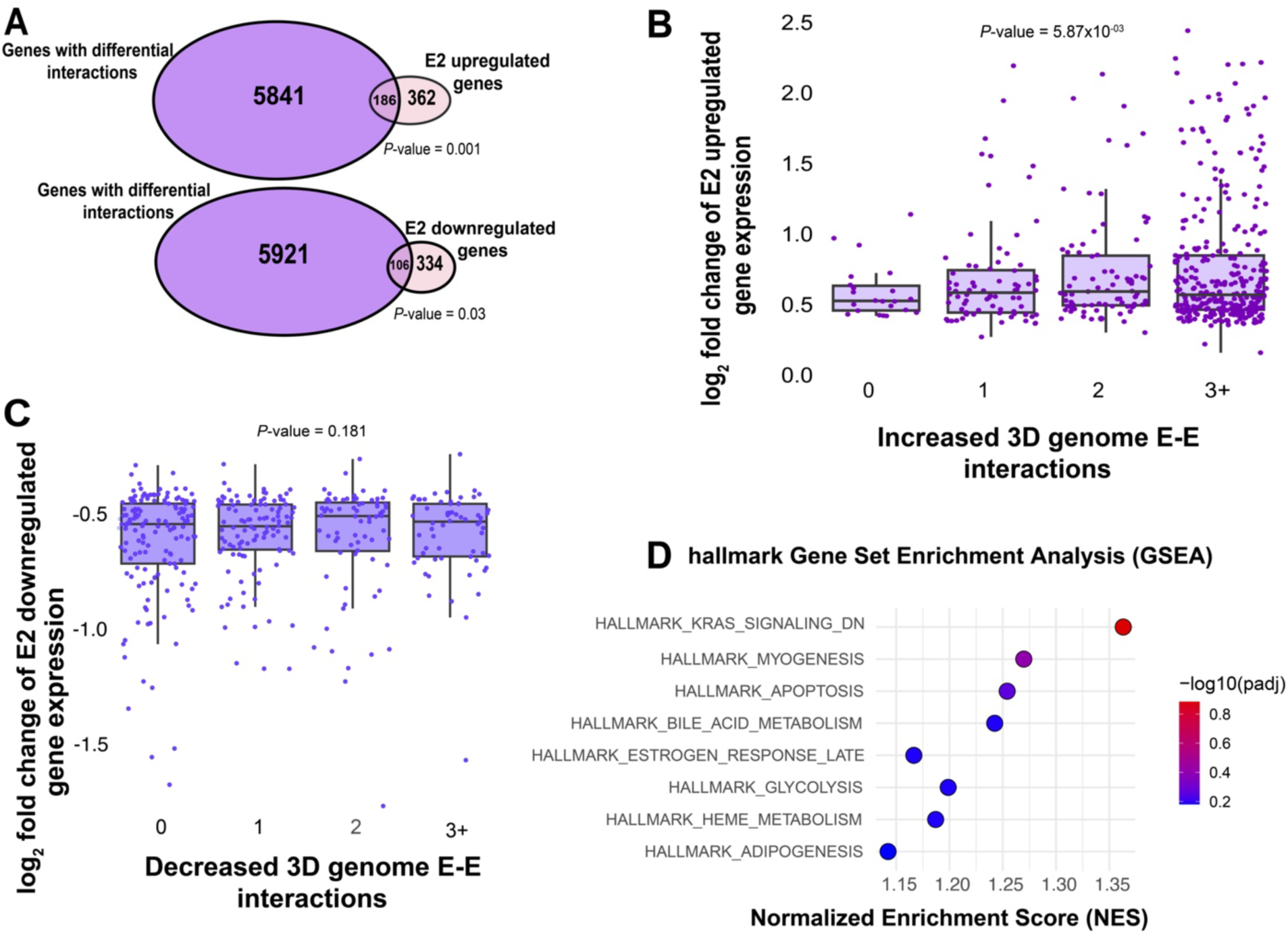
3D genome structure changes upon estrogen treatment associate with differential gene expression. A) Overlap is shown between E2 upregulated or downregulated genes (purple) and all genes involved with differential 3D interactions (pink) in Ishikawa cells. (B-C) Boxplots demonstrate relative expression of E2 upregulated (B) and downregulated (C) genes associated with increasing (B) or decreasing (C) 3D genome interactions in Ishikawa cells, respectively. *P*-values are from t-test on the linear regression coefficient being different from 0. D) The GSEA rank of hallmark gene set enrichments based on the number of 3D genome interactions within 100 kb of each gene’s TSS in Ishikawa cells is shown.

We next determined whether differential 3D genome interactions were associated with larger transcriptional responses to E2. Using linear regression models, we found that more differential Enhancer–enhancer interactions were associated with larger upregulated (Figure 2B; *P*-value =0.00587, t-test on coefficient) but not downregulated (Figure 2C; *P*-value = 0.181, t-test on coefficient) transcriptional changes to E2 treatment in Ishikawa cells; however, HCI-EC-23 did not show a significant association with either upregulation (Figure S2B; *P*-value = 0.366, t-test on coefficient) or downregulation (Figure S2C; *P*-value = 0.706, t-test on coefficient) in response to E2 treatment. We did not find a significant association between the number of differential loops and basal expression levels, indicating that the connection between the E2 driven transcriptional changes and the number of differential loops in Ishikawa cells is not caused by differences in basal expression levels. Gene Set Enrichment analysis (GSEA) indicated no significant pathways distinguishing genes associated with increased or decreased 3D genome interactions in response to E2 treatment, in both Ishikawa (Figure 2D) and HCI-EC-23 cells (Figure S2D); however, estrogen response was one of the top hallmark gene sets based on enrichment score in Ishikawa cells. Overall, these results suggest that chromatin looping changes in response to estrogen treatment are associated with differentially expressed genes in Ishikawa cells, but to a lesser extent in HCI-EC-23, hence most subsequent studies involving E2 response were conducted in Ishikawa cells. Additionally, these results reveal several genes that exhibit 3D genome interaction changes or gene expression changes, but not both.

### ER binding events are enriched in differential 3D genome interactions

Since ER is the main mediator of the transcriptional response to estrogen in endometrial cancer cells, we next evaluated the relationship between ER genomic binding and chromatin looping. Integration of previously collected ER ChIP-seq data (Rush et al. 2022; Gertz et al. 2013) with the HiChIP data revealed that the majority of ERBS were found in 3D genome interactions (69% in Ishikawa and 70% in HCI-EC-23) (Figure 3A). Approximately 22% of ERBS did not harbor H3K27ac; therefore, 3D genome interactions involving these sites might have been missed for technical reasons, while less than 10% of ERBS that had detectable H3K27ac signal were not found in detectable 3D genome interactions (Figure 3A).

**Figure 3.**
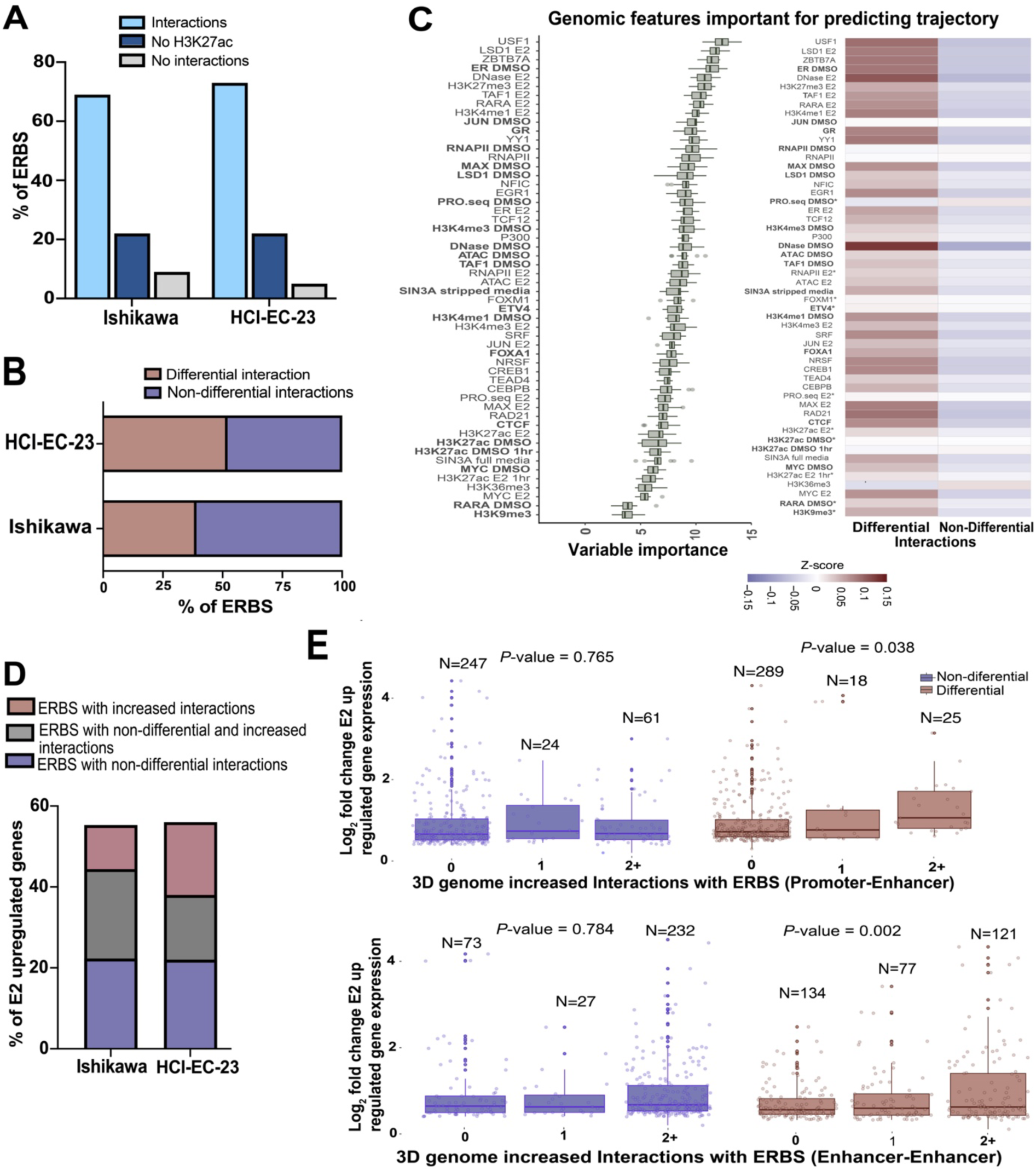
ERBS exhibit distinctive features when involved in differential 3D genome interactions. (A) The bar graph portrays all ERBS associated with 3D genome interactions and their classifications in endometrial cell lines. (B) Of the ERBS found in 3D genome interactions (light blue in panel A), the bar graph indicates the percentage that anchors differential 3D genome interactions. (C, left) A ranking of genomic features based on their importance for ERBS associated with differential or non-differential 3D genome interactions (exhibited in panel B) is shown. Datasets shown in bold were performed in the absence of ER activation. (C, right) Heatmap displays the average signal intensity for top-ranked genomic features clustered by ERBS with differential or non-differential 3D genome interactions. Each feature’s distributions were significantly different between groups based on a Kruskal Wallis test (False Discovery Rate < 0.05) with the exception of features marked with an *. (D) Bar graph shows the percentage of E2 upregulated genes associated with ERBS in differential or non-differential 3D genome interactions (from panel B) within 100 kb of their TSS. (E) Box plots show fold change for E2 upregulated genes with at least one ERBS within 100 kb of their TSS in Ishikawa cells split by the number of non-differential (left, blue) or differential (right, red) Promoter–enhancer (top) or Enhancer–enhancer (bottom) chromatin loops anchored by ERBS. *P*-values are calculated as t-tests of the linear regression coefficients being different from 0.

We next evaluated ERBS associated with changes in 3D genome interactions caused by E2 treatment. We observed a very strong enrichment of ERBS in differential 3D genome interactions (Figure 3B, *P*-value = 2.46x10^-37^, hypergeometric distribution), with nearly half of ERBS serving as anchors for differential 3D genome interactions. The majority of these ERBS (71.9 % in Ishikawa and 79.2 % in HCI-EC-23) were associated with increased 3D genome interactions as opposed to decreased 3D genome interactions. To understand the differences between ERBS associated with differential or non-differential 3D genome interactions, we used Boruta, a random forest feature selection approach, to look for genomic features in Ishikawa cells that distinguish ERBS anchoring differential 3D genome interactions (Ginley-Hidinger et al. 2024). Several transcription factors as well as chromatin accessibility features were enriched at ERBS with differential 3D genome interactions, while ERBS associated with non-differential 3D genome interactions exhibited features of actively transcribed region, particularly in the absence of E2, including PRO-seq signal and H3K36me3 (Figure 3C). These results suggest that ERBS associated with non-differential interactions are already engaged in transcription prior to E2 treatment, while ERBS associated with differential 3D genome interactions are open regulatory regions with less active transcription; however, the effect sizes are relatively small indicating that these features represent only part of the explanation.

To determine how ERBS-associated changes in 3D genome interactions relate to the E2 transcriptional response, we first identified ERBS within 100 kb of the TSS of E2 upregulated genes in Ishikawa cells and then split ERBS based on whether they served as anchors for differential, non-differential, or both types of 3D genome interactions. We observed that ERBS involved in differential and non-differential 3D genome interactions were similarly enriched for E2 upregulated genes (Figure 3D). In addition, ERBS with differential or non-differential interactions were equally likely to regulate genes that respond early or late during an 8-hour time course (Ginley-Hidinger et al. 2024) (*P*-value = 0.55, Fisher’s exact test). Next, we used linear regression to determine if ERBS associated with differential 3D genome interactions were predictive of larger transcriptional responses to E2. We observed that the size of the transcriptional response to E2 was positively correlated with the number of Promoter–enhancer or enhancer–enhancer differential 3D genome interactions anchored by ERBS, but not with the number of Promoter– enhancer or enhancer–enhancer non-differential 3D genome interactions anchored by ERBS (Figure 3E). It is important to point out that the connections between ERBS and target genes exhibited multiple levels of redundancy; 24% of target genes were connected to more than one ERBS through Promoter–enhancer interactions. In addition, 49% of target gene promoters were serially connected to two ERBS (i.e., an ERBS interacts with another ERBS that anchors a Promoter–enhancer interaction). Overall, our findings indicate that ER genomic binding sites are equally likely to be involved with non-differential or differential chromatin loops and that ERBS in differential 3D interactions are more open and less transcriptionally active prior to E2 treatment, but are associated with larger changes in the expression of their target genes upon E2 induction.

While differential 3D genome interactions were highly enriched for ERBS upon E2 treatment, most differential 3D genome interactions were not associated with ERBS in Ishikawa (89%) and HCI-EC-23 (90%) cells. To identify the features of differential 3D genome interactions that are not anchored by ERBS, we first classified non-differential or differential 3D genome interactions in both cell lines based on whether a chromatin loop anchor overlapped an ERBS (ER primary interaction) or whether a chromatin loop anchor interacted with an ERBS through another chromatin loop (ER secondary interaction); the remaining interactions were categorized as ER-unrelated interactions (Figure S3A and S3B). We used Boruta analysis to identify features that distinguish ER-unrelated differential 3D genome interactions from ER-unrelated non-differential 3D genome interactions in Ishikawa cells. The results showed that ER-unrelated differential 3D genome interactions were enriched for several transcription factors and chromatin accessibility (Figure S3C). When we further categorized ER-unrelated differential 3D genome interactions based on whether they increased or decreased, we found that decreased ER-unrelated 3D genome interactions exhibited a stronger enrichment of transcription factors than increased 3D genome interactions; however, the effect sizes are relatively small. On the contrary, increased 3D genome interactions without ERBS were uniquely associated with features of active transcription, including PRO-seq signal and RNA Polymerase II (Figure S3D). While the question remains as to why thousands of non-ER associated chromatin loops are changing in interaction frequency, the strongest pattern that we observed is a reduction in chromatin looping at anchors with a high level of transcription factor binding and chromatin accessibility, which may indicate competition for activating regulatory events when ER is active.

### ERBS in differential chromatin loops are slightly more likely to regulate transcription

The analysis described above led us to hypothesize that ERBS anchoring differential 3D genome interactions upon E2 treatment are more likely to impact target gene transcription. To investigate the functional importance of different types of ERBS, we used the CRISPR-based Enhancer-i system (Carleton et al. 2018) to repress the activity of individual ERBS and measure the gene expression response of target genes to an 8-hour E2 treatment. We first classified ERBS into differential or non-differential 3D genome interactions and further classified the sites into Promoter–enhancer or Enhancer–enhancer interactions, targeting at least 5 ERBS in each category. Results from our previously published Enhancer-i study (Carleton et al. 2017) were combined with experiments performed for this study (Table S1).

In the Promoter–enhancer category, 80% of ERBS with increased differential 3D genome interactions and 67% of ERBS with non-differential 3D genome interactions were found to be functionally important for the E2 transcriptional response (Figures 4A and S4A). In the Enhancer– enhancer category, 86% of ERBS with increased differential 3D genome interactions and 67% of ERBS with only non-differential 3D genome interactions were found to be functional (Figure 4C and S4A). For example, there are 3 ERBS near *GDPD5*, an E2 upregulated gene. ERBS 1, which anchors an increased Promoter–enhancer interaction, significantly impacts the expression of *GDPD5* (Figure 4B). ERBS 2 and 3 near *GDPD5* anchor non-differential Enhancer–enhancer interactions and significantly impact *GDPD5* expression (Figure 4B). Another example is the 4 ERBS near *TACSTD2*, an E2 upregulated gene; ERBS 1, 2, and 4 anchor increased Enhancer– enhancer interactions, while ERBS 3 anchors non-differential Enhancer–enhancer interactions. Targeting ERBS 2 or 4 significantly impacted the expression of *TACSTD2*, whereas targeting ERBS 1 or 3 did not significantly impact *TACSTD2* expression (Figure 4D). The successful targeting of each ERBS by Enhancer-i was validated using ChIP-seq (Figure S5A). Overall, 83% of ERBS in differentially increased 3D genome interactions, and 67% of ERBS in non-differential 3D genome interactions demonstrated a significant impact on the E2 transcriptional response; however, this difference was not significant with the 24 ERBS tested (*P*-value =0.4995; Fisher’s exact test). In addition, we examined the transcriptional effect of each ERBS and found that ERBS that anchor differentially increased 3D genome interactions reduced the E2 response by an average of 62% whereas ERBS that do not anchor differential interactions lead to an average reduction of 48% when targeted by Enhancer-i (Figure 4E), which was not statically significant (*P*-value = 0.5037, t-test). These results suggest that ERBS associated with increased 3D genome interactions trend toward a higher likelihood of being functional and causing larger effects on transcription than ERBS associated with non-differential 3D genome interactions. However, the difference is subtle, and several ERBS that only anchor non-differential chromatin loops contribute to the transcriptional response to estrogen.

**Figure 4.**
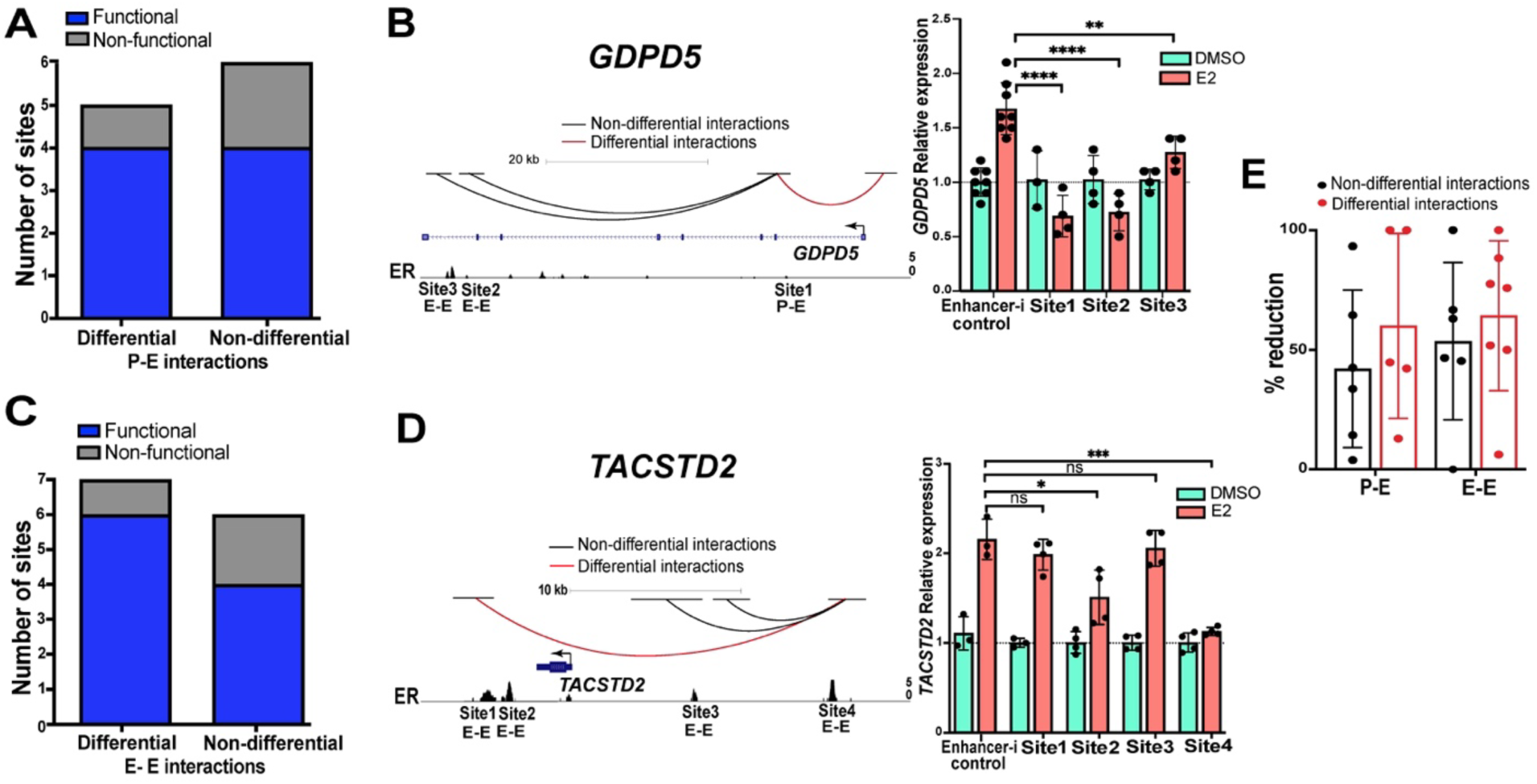
ERBS in differential chromatin loops are subtly more important for the transcriptional response to estrogen. (A/C) The functionality of ERBS based on Enhancer-i that are anchors of differential and non-differential Promoter–enhancer (P-E; A) or Enhancer–enhancer (E-E; C) interactions is shown. (B/D, left) Genome browser tracks of *GDPD5* (B) and *TACSTD2* (D) show differential (red) and non-differential (black) 3D genome interactions as well as ER ChIP-seq signal. (B/D, right) Relative expression of *GDPD5* (B) and *TACSTD2* (D) normalized to DMSO when a candidate ERBS is targeted with Enhancer-i upon DMSO (green) or E2 (orange) treatment is displayed. Error bars represent the SD; ****p < 0.00001, ***p < 0.001, **p < 0.01 and *p < 0.05, unpaired t-test, while ns depicts statistical insignificance. (E) The average percent reduction is shown for each ERBS targeted with Enhancer-i split by the different types of 3D interactions. Each dot represents the average percent reduction for one ERBS, and error bars represent SD.

### E2 upregulated gene promoters are likely to reside in physical hubs that can cooperate in an estrogen response

In addition to Promoter–enhancer and Enhancer–enhancer interactions discussed above, our analysis revealed several Promoter–promoter interactions in the HiChIP data. When analyzing the promoters of estrogen-induced genes, we initially looked at promoter proximity in the linear genome. Evaluation of the distance between promoters showed that E2 upregulated genes cluster together in the genome more than expected by chance (*P*-value = 0.00031, Wilcoxon rank sum test) (Figure 5A), indicating that some promoters may be coregulated by E2. Using a 150 kb distance cutoff for potentially coregulated gene pairs, we identified 81 E2 upregulated and 49 E2 downregulated gene pairs. Comparison to HiChIP data revealed that many of the paired E2 regulated gene promoters physically interacted with each other, where more promoter interactions were observed within E2 upregulated gene pairs, both with and without E2 treatment, compared to randomly selected gene pairs within 150 kb (Figure 5B). These observations suggest that many E2 upregulated genes reside in pre-existing promoter interaction hubs, which raises an interesting question as to whether a functional relationship exists between these interacting promoters in terms of their transcriptional response to estrogen.

**Figure 5.**
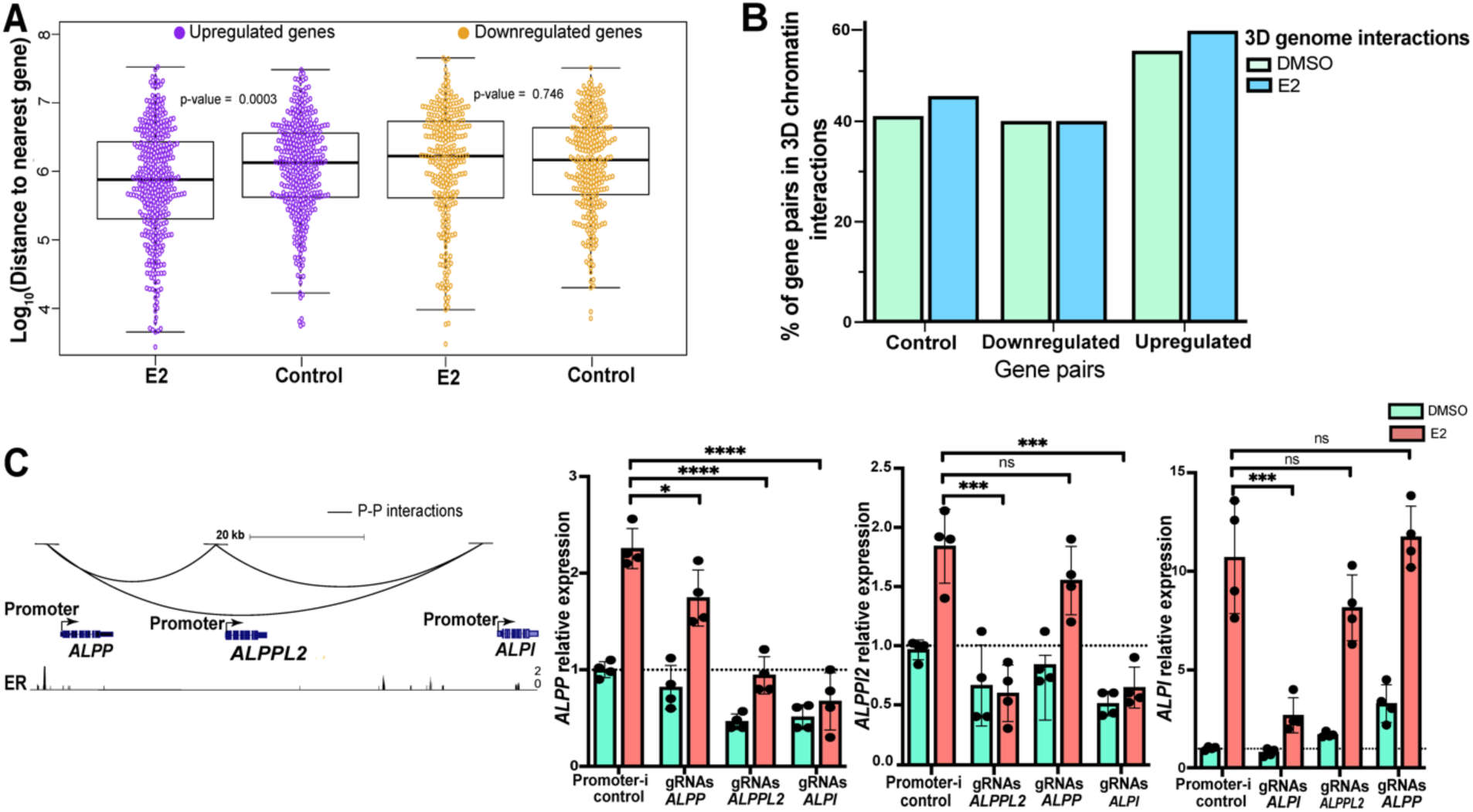
E2 upregulated gene promoters are enriched in physical interactions that can mediate their transcriptional response to estrogen. (A) The plot displays the genomic distances among E2 responsive genes or randomly selected control genes that match in the number of genes. Upregulated genes and matching control genes are shown in purple; downregulated genes and matching controls are shown in yellow. (B) Bar plot shows the percentage of E2 regulated or randomly selected (control) gene pairs within 150 kb that have Promoter–promoter interactions (left, control genes; middle, downregulated genes; right, upregulated genes) observed under DMSO (green) or E2 (blue) treatment conditions. (C, left) Genome browser tracks of *ALPP-ALPPL2-ALPI* gene cluster displaying Promoter–promoter interactions and ER ChIP-seq signal. (C, right) Relative expression with DMSO (green) or E2 (orange) treatment of each gene when the genes’ promoters in the cluster are targeted by Enhancer-i is displayed. Error bars represent the SD; ****p < 0.00001, ***p < 0.001, **p < 0.01 and *p < 0.05, unpaired t-test, while ns depicts statistical insignificance.

To explore the functional relationship between interacting promoters, we targeted the promoters of an E2 upregulated gene cluster (*ALPP-ALPPL2-ALPI*) using Enhancer-i and determined how the activity of one gene in the cluster impacted other cluster genes. The *ALPP-ALPPL2-ALPI* gene cluster exhibited cooperative gene expression, where targeting a single promoter affected not only the targeted gene’s expression but also the expression of other genes in the cluster (Figure 5C). However, the cooperation was not equal; *ALPI* had the most important promoter in the group as it impacted all genes, *ALPPL2*’s promoter cooperated with *ALPP*, and *ALPP*’s promoter was only important for its own expression. The successful targeting of each promoter by Enhancer-i was validated using ChIP-seq (Figure S5B). Overall, these results revealed that E2 upregulated gene promoters can reside in physical hubs and potentially cooperate in the transcriptional response to estrogen.

## Discussion

The relationship between 3D genomic structure and gene expression is complex and has been described as context-specific by several studies (Pollex et al. 2024b; Chen et al. 2024). Some studies have reported minimal changes in expression when large perturbations in genome architecture are observed, while other studies have found large changes in expression for particular genes when the genome structure is altered. In this study, we functionally dissected the relationship between changes in 3D genome structure and gene regulation in the context of estrogen signaling in endometrial cancer cells. Our focus is on estrogen signaling for several reasons, including endogenous activation of ER by estrogen in endometrial cancer cells, ER activation through estrogen signaling is a key oncogenic event in endometrial cancer (Rodriguez et al. 2019), and ER is a well-studied transcription factor, which provides a strong foundation for interpretation of results. We found that 1 hour of estrogen treatment was sufficient to significantly impact tens of thousands of 3D genome interactions. The observed changes were enriched at genes that change expression; however, many genes harbor differential chromatin loops without concomitant changes in gene expression. This observation could be caused by changes in 3D interactions that are unrelated or at least nonproductive concerning gene regulation, or the non-responsive genes could be unresponsive to an 8-hour E2 treatment and may require other stimuli or a different amount of treatment time to respond.

Since ER is the main mediator of transcriptional effects of estrogen in these endometrial cancer models, we analyzed 3D genomic interactions in the context of ER genomic binding. Most ERBS served as anchors for chromatin loops, and approximately half of ERBS anchored chromatin loops that increase in frequency when cells are treated with estrogen for 1 hour. Further analysis uncovered a pattern where ERBS with differential 3D genome interactions have more accessible chromatin, several transcription factors bound, and less transcription activity based on PRO-seq and RNA polymerase II binding. This observation is consistent with sites of active transcription exhibiting higher motility (Gu et al. 2018) and findings in breast cancer cells that estrogen-dependent enhancers increase contact frequencies with their promoters in a transcription-independent manner (Gomez Acuna et al. 2024). Our findings, together with previous studies, suggest a model in which ER binding does not typically impact 3D genome interaction frequencies at regulatory regions that are already transcriptionally active and already highly motile, while ER binding to transcriptionally inactive sites with accessible chromatin is more likely to increase chromatin looping frequencies. While this is an interesting model, it is unlikely to be the complete explanation since these features are only subtly different between ERBS that anchor differential and non-differential 3D genome interactions. In addition, no transcription factor binding motifs were significanlty enriched between ERBS that anchor differential loops and ERBS that anchor only non-differential loops.

Despite a strong enrichment of ERBS at differential chromatin loops, the majority of differential 3D genome interactions were not connected to ER in both cell lines. Further analysis uncovered that ER-unrelated differential 3D genome interactions were anchored by several transcription factors and more open chromatin than ER-unrelated non-differential 3D genome interactions. Most decreased 3D genome interactions were linked to several transcription factors other than ER and associated with less transcription, while the increased 3D interactions without ER were associated with active transcription. These results were consistent with previous ER-positive breast cancer findings, where differential 3D genome interactions that changed upon estrogen were predominantly associated with transcription factors, such as CTCF (Luo et al. 2020; Witcher and Emerson 2009), which was similarly enriched with decreased ER-unrelated 3D genome interactions in this study. Overall, these findings revealed that changes in 3D genome interactions upon E2 treatment are potentially the result of a combined effort or competition between ER and several other genomic features, yet further studies are needed to understand the mechanisms underlying these ER-unrelated changes.

Through correlative analysis, we found that increases in 3D genome interaction frequency were associated with larger transcriptional responses to estrogen. This finding led us to hypothesize that ERBS with changes in chromatin looping frequency have larger expression effects on their target genes. To test this hypothesis, we took a functional approach by targeting different sets of ERBS with Enhancer-i. In total, we analyzed 24 ERBS split into groups of differential or non-differential 3D interactions as well as Promoter–enhancer or Enhancer– enhancer interactions. The overall patterns that we observed were very similar between ERBS involved in Promoter–enhancer and Enhancer–enhancer interactions, suggesting that Enhancer– enhancer 3D contacts are likely important for the ability of ERBS to regulate gene expression. Across both types of contacts, we found that ERBS that anchor differential 3D genome interactions functionally impacted the transcriptional response to estrogen at a higher percentage than ERBS that only anchor constant 3D interactions. However, this difference was not significant, possibly due to our study being underpowered to detect a subtle difference in functionality between these types of ERBS. The observed subtle difference in functionally between ERBS that anchor differential or non-differential 3D genome interactions may explain why we observed only a modest overlap between genes with differential 3D genome interactions and genes that change expression upon E2 treatment (Figure 2A). While differential chromatin looping upon E2 treatment may lead to larger changes in the transcriptional response, it is clear that ERBS without changes in 3D genome interactions play an important role in transcription regulation.

In addition to chromatin loops that involve enhancers, we also observed several Promoter–promoter 3D genome interactions between genes that are upregulated by estrogen. This finding goes along with our observation that estrogen upregulated genes cluster in close proximity within the genome more than expected by chance. Previous studies on chromatin interactions associated with RNA polymerase II have revealed that most genes with Promoter– promoter interactions are transcribed cooperatively (Pollex et al. 2024a; Li et al. 2012; Xu et al. 2011), while other findings have detected transcriptional competition among neighboring genes (Cho et al. 2018). To determine how 3D connected promoters of estrogen upregulated genes impact one another, we used Enhancer-i to target a gene’s promoter and analyzed how other genes with connected promoters respond to estrogen. A cooperative relationship was observed among the three E2 upregulated genes, whose promoters physically interacted in 3D space; however, the promoter strength was not equal among these genes. This study only surveyed 3D genome interactions of one group of estrogen regulated genes, therefore competition and other complex relationships between the promoters of estrogen regulated genes may exist within different gene clusters.

## Methods

### Cell culture

Ishikawa and HCI-EC-23 cells were grown in full media: RPMI 1640 medium (Thermo Fisher Scientific) supplemented with 10% fetal bovine serum (FBS) (Thermo Fisher Scientific) and 1% penicillin-streptomycin (Thermo Fisher Scientific) and were incubated at 37°C with 5% CO_2_. Cells were transferred and maintained in hormone-depleted media consisting of phenol red-free RPMI 1640 (Thermo Fisher Scientific) supplemented with 10% charcoal-dextran treated FBS (Sigma Aldrich) and 1% penicillin-streptomycin (Thermo Fisher Scientific) for 5 days prior to E2 treatment.

### HiChIP

The 3D genome structure was analyzed using the published HiChIP protocol (Mumbach et al. 2017), which consists of three main phases, including in situ contact generation, sonication, and chromatin immunoprecipitation. In brief, crosslinked nuclei were extracted from 10 million cells treated with either 10 nM E2 or DMSO for 1 hour and digested with the restriction enzyme DpnII. The restriction fragment overhangs were filled with biotinylated dATP to mark the DNA ends. Biotin-marked DNA ends were ligated to covalently link interacting regions. Following proximity ligation, the nuclei with *in situ* generated contacts were pelleted and suspended in a nuclear lysis buffer. Crosslinked chromatin was sonicated using an EpiShear probe-in sonicator (Active Motif) with three cycles of 30 seconds at an amplitude of 40% with 30 seconds of rest between cycles. The antibody that recognized H3K27ac (Abcam, ab4729) was used to pull down crosslinked chromatin associated with H3K27ac overnight at 4C. Protein A/G magnetic beads (Invitrogen) were used for crosslinked chromatin purification. Upon reverse crosslinking, biotin-labeled post-immunoprecipitation DNA was captured by streptavidin C1 magnetic beads (Invitrogen). Tn5 (Illumina) was used to fragment captured biotin-labeled DNA and prepare molecules for PCR generation of sequencing libraries.

HiChIP libraries were sequenced on an Illumina NovaSeq 6000 as paired-end 50 basepair reads to an average depth of 300–400 million read-pairs per sample. Reads were aligned to the human hg19 reference genome using HiC-Pro (Servant et al. 2015) to extract informative unique paired-end tags (PETs). The hg19 build of the human genome was used for all genomic analyses. We do not believe that realigning reads to the GRCh38 genome build would substantially change results, because we are restricting our analyses to uniquely alignable regions of the genome. Hichipper (Lareau and Aryee 2018) was used to perform restriction site bias-aware modeling of the output from HiC-Pro and to call chromatin loops. These chromatin loops were filtered where chromatin loops with at least one or more reads in both biological replicates of a particular treatment (E2 or DMSO) were kept and used for downstream analyses. Chromatin interactions that significantly change upon 1 hour of E2 induction were identified using DESeq2 (Love et al. 2014) with an adjusted *P*-value cutoff of 0.05. Enhancer–enhancer chromatin loops were assigned to genes using GREAT (McLean et al. 2010) with a cutoff of 100 kb. Promoters were defined as any anchor that overlapped within 500bp of the transcription start sites of University of California Santa Cruz (UCSC) Known Genes.

### Enhancer interference

Guide RNAs (gRNAs) targeting regions of interest for Enhancer-i (4 gRNAs per targeted region) were designed using the Benchling gRNA design tool (see Table S2 for sequences). gRNAs were cloned into pGL3-U6-PGK-Puro (Addgene 51133, a gift from Xingxu Huang) as previously described (Carleton et al. 2017). Ishikawa cells expressing SID4x-dCas9-KRAB fusion protein (Carleton et al. 2017) were cultured in full media supplemented with 600 ng/uL G418 and incubated at 37°C with 5% CO_2_. Six days before transfection, cells were transferred and maintained in hormone-depleted media with 600ng/uL G418. 24 hours before transfection, cells were plated in a 24-well plate at 90,000 cells per well and transfected with gRNAs using FuGENE HD (Promega) at a manufacturer-suggested 3:1 reagent:DNA ratio. At 36 hours post-transfection, cells were treated with 1 μg/mL puromycin to select for cells successfully transfected with gRNA plasmids. Selected cells were induced 24 hours later with 10nM E2 or DMSO (vehicle) for 8 hours. Significance was determined using unpaired t-tests. Normality was evaluated for each site and for the control of each gene using a Shapiro-Wilk test and results were found to be consistent with a normal distribution in all cases.

### ChIP-seq

Ishikawa cells expressing the SID4x-dCas9-KRAB fusion protein were cultured as described in the above section. 24 hours before transfection, cells were plated in 15 cm dishes at approximately 60% confluency. gRNAs targeting 24 different sites were combined into four separate pools; each pool contained gRNAs that targeted 6 different sites nearby different genes and were transfected into plated cells. The remaining steps, including E2 treatment, were conducted as described in the above section. ChIP-seq libraries were generated as previously described (Reddy et al. 2009) and the antibody used for this study was FLAG M2 (Sigma-Aldrich F1894) that targeted a FLAG tag on the SID4x-dCas9-KRAB fusion. The ChIP-seq libraries were sequenced on an Illumina NovaSeq X as paired-end 150 bp. Bowtie (Langmead et al. 2009) was used to trim the reads to 50 bp by removing 100bp from the 3’ ends and aligning the trimmed reads to the human hg19 reference genome. SAMtools view (Li et al. 2009) was used to convert aligned sequences to BAM format; MACS2 (Zhang et al. 2008) was used to create bedgraphs. Bedgraphs were converted to bigWig using UCSC’s BedgraphToBigWig tool and visualized on the UCSC Genome Browser.

### RNA isolation and qRT-PCR

Cells were lysed in RLT Plus (Qiagen) supplemented with 1% beta-mercaptoethanol (Sigma). RNA was extracted and purified using the ZR-96-well Quick-RNA kit (Zymo Research). Gene expression was quantified using reverse transcription quantitative PCR (RT-qPCR) at 40 cycles on a CFX Connect light cycler (BioRad). qPCR reaction mixtures were set up with reagents from the Power SYBR Green RNA-to-Ct 1-step kit (Thermo Fisher Scientific) with 25-50 ng RNA per reaction and primers are listed in Table S3. Melt curve analysis was conducted to assess amplicon qPCR length and specificity. Relative expression was determined using the ΔΔCt method; the reference gene used for this assay was *CTCF*. The significance of gene expression change between experimental conditions was determined by unpaired t-test with a *P*-value cutoff of 0.05.

### Analysis of important features

The Boruta analysis was performed as described by Ginley-Hidinger et al. (Ginley-Hidinger et al. 2024) and information on the datasets used can be found in Table S4. For the analysis looking at differences between differential and non-differential chromatin loops, each anchor was included separately. Due to disparities in the size of different sets, we down-sampled the larger set and Boruta (Kursa and Rudnicki 2010) was run using 100 random downsamplings. For linear regression analysis of the association between the numbers of chromatin loops and expression responses to E2, data was first compiled in a gene-centric table with each row containing a gene, the log_2_ expression fold change after E2 treatment, and the sum of chromatin loops that interact with that gene in Enhancer–enhancer, Promoter–enhancer, and Promoter– promoter interactions. We used the lm function in R version 4.3.1 (Team 2024) as follows: lm(formula = Log_2_ Fold Change ∼ Enhancer–enhancer + Promoter–enhancer + Promoter– promoter). This analysis was performed using all chromatin loops or only ERBS anchored chromatin loops. Significance was determined using a t-test on the coefficients with a cutoff of 0.05. The GSEA analysis was performed using the number of differential chromatin loops where decreased loops were subtracted from increased loops within 100 kb of the TSS for ranking genes.

## Data access

All raw and processed sequencing data generated in this study have been submitted to the NCBI Gene Expression Omnibus (GEO; https://www.ncbi.nlm.nih.gov/geo/) under accession number GSE269670.

## Acknowledgments

Funding for this work came from the National Institutes of Health (NIH)/National Human Genome Research Institute (NHGRI) R01 HG008974 to J.G., the American Association of University Women to H.A., and the Huntsman Cancer Institute. Research reported in this publication utilized the High-Throughput Genomics Shared Resource at the University of Utah and was supported by NIH/National Cancer Institute (NCI) award P30 CA042014. We thank Jake Polaski for experimental and writing guidance and Gertz lab members for their suggestions on the study and manuscript.

## Author contributions

Conceptualization, H.A. and J.G.; Methodology, H.A., X.Y., and J.G.; Investigation, H.A., C.M.R, and N.K.; Formal Analysis, H.A., A.R., J.M.V., M.G., and J.G.; Writing - Original Draft, H.A. and J.G.; Writing, Review & Editing, all authors; Supervision, J.G., and X.Z.; Funding Acquisition, H.A. and J.G.

