## Supplementary figures and tables for "Estrogen-induced chromatin looping changes identify a subset of functional regulatory elements"

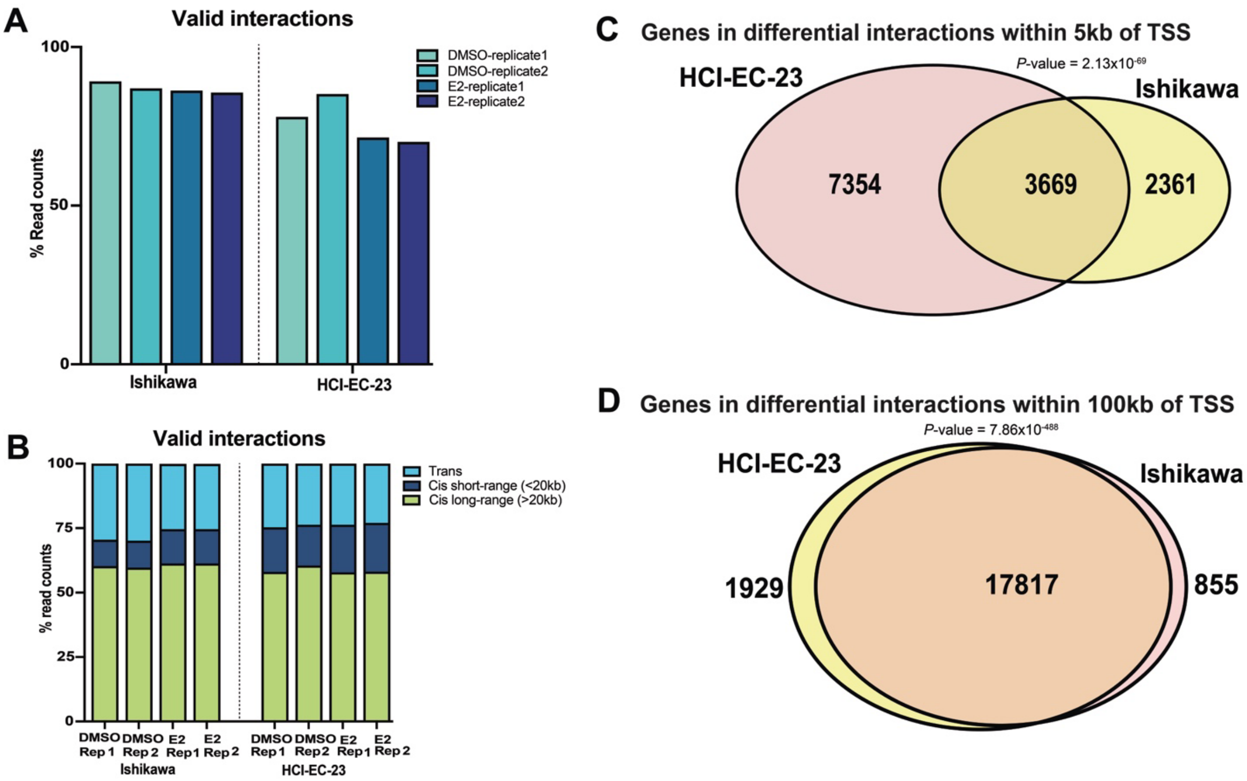


**Supplementary Figure 1 (related to Figures 1 and 2). Quality control plots for HiChIP and diagram of genes with differential 3D interactions**. A) Bar plot displays percentage of valid HiChIP reads in Ishikawa (left) and HCI-EC-23 (right) sample replicates. B) The distribution of trans (light blue), cis short-range (dark blue), or cis long-ranger (green) valid interactions in Ishikawa (left) and HCI-EC-23 (right) sample replicates is shown. (C-D) Venn diagrams display the overlap of genes associated with differential 3D interactions that have anchors within 5kb of TSS (C) or within 100kb of TSS (D) between HCI-EC-23 (yellow) and Ishikawa (pink).

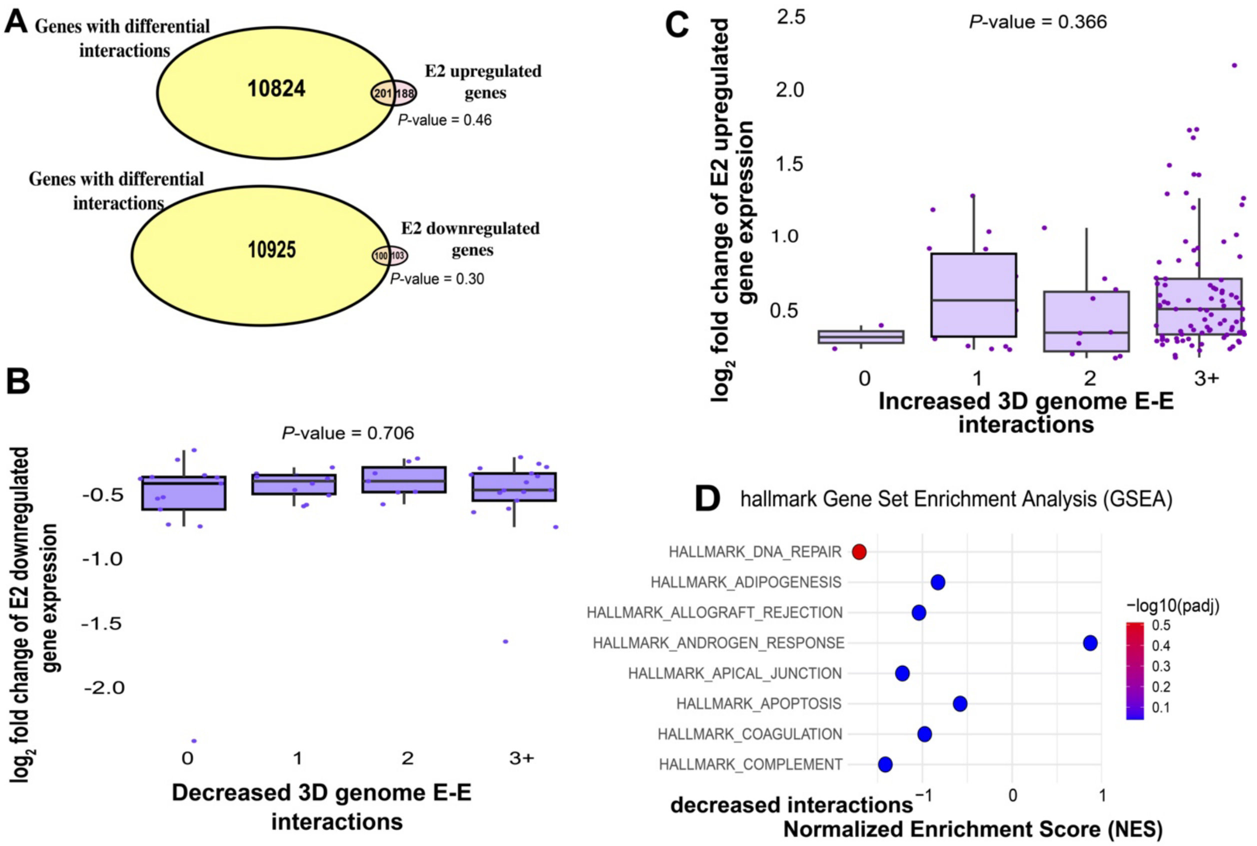


**Supplementary Figure 2 (related to Figure 2). Comparison of genes associated with differential interactions and estrogen regulated genes**. A) Overlap reveals E2 upregulated or downregulated genes (pink) that exhibit differential 3D interactions in HCI-EC-23 cells. (B-C) Boxplots demonstrate relative expression of E2 upregulated (B) and downregulated (C) genes associated with increasing (B) or decreasing (C) 3D genome interactions in HCI-EC-23 cells, respectively. P-values are from t-test on the linear regression coefficient . D) The GSEA rank of hallmark gene set enrichments based on the number of 3D genome interactions within 100 kb of each gene's TSS in HCI-EC-23 cells is shown.

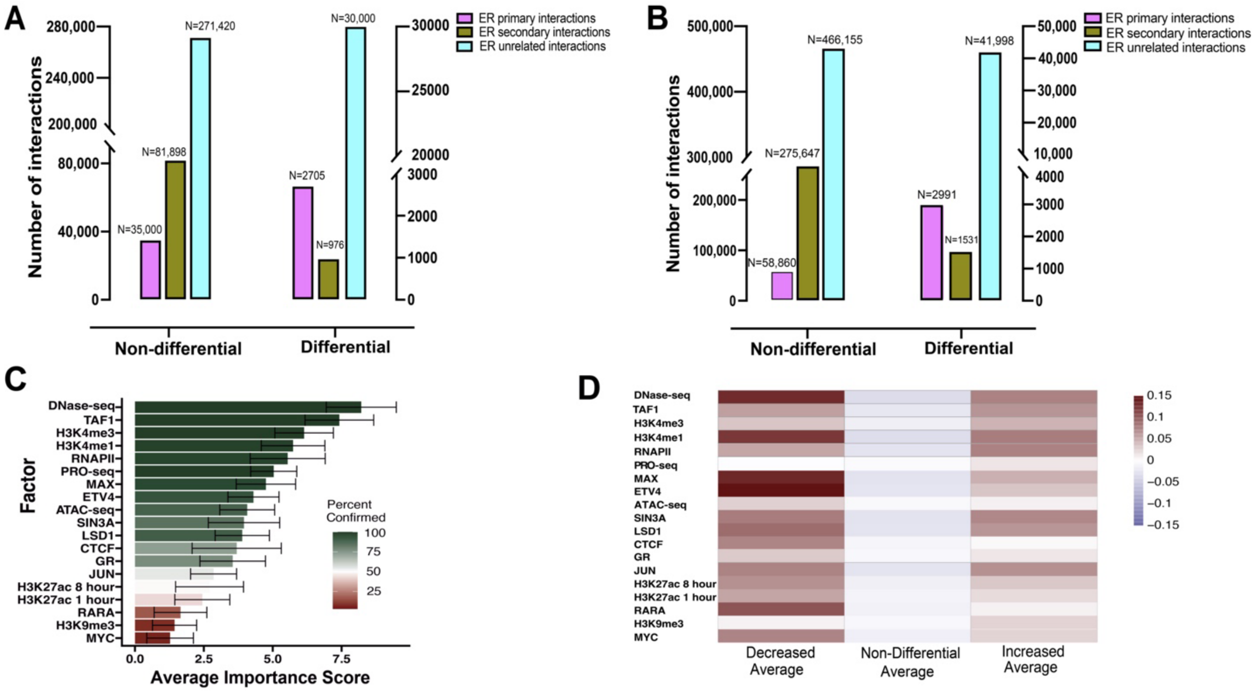


**Supplementary Figure 3 (related to Figure 3). Characterization of 3D genome interactions associated with ER binding events. (**A-B) The distribution of non-differential (left) or differential (right) 3D interaction anchors based on whether an anchor is an ERBS (magenta), an anchor interacts with an anchor that is an ERBS (green), or the two anchors of an interaction aren't involved in any loops that include an ERBS as an anchor (blue) is shown for Ishikawa (A) or HCI-EC-23 (B); N represents the number of 3D interactions for each category. C) The Boruta importance scores are shown for the analysis of anchors in differential compared to non-differential loops for ER-unrelated interactions. Only datasets collected in the absence of ER activation are shown. Error bars represent SD from 100 random down-sampled runs, and colors represent the percentage of time a variable is confirmed as important. D) Heatmap displays the average signal intensity for top-ranked genomic features from a Boruta analysis comparing ER-unrelated loops that are reduced, non-differential, or increased with E2 treatment in Ishikawa cells. Only datasets collected in the absence of ER activation are shown. Each feature's distributions were significantly different between at least one pair of groups based on a Kruskal Wallis test (False Discovery Rate < 0.05).

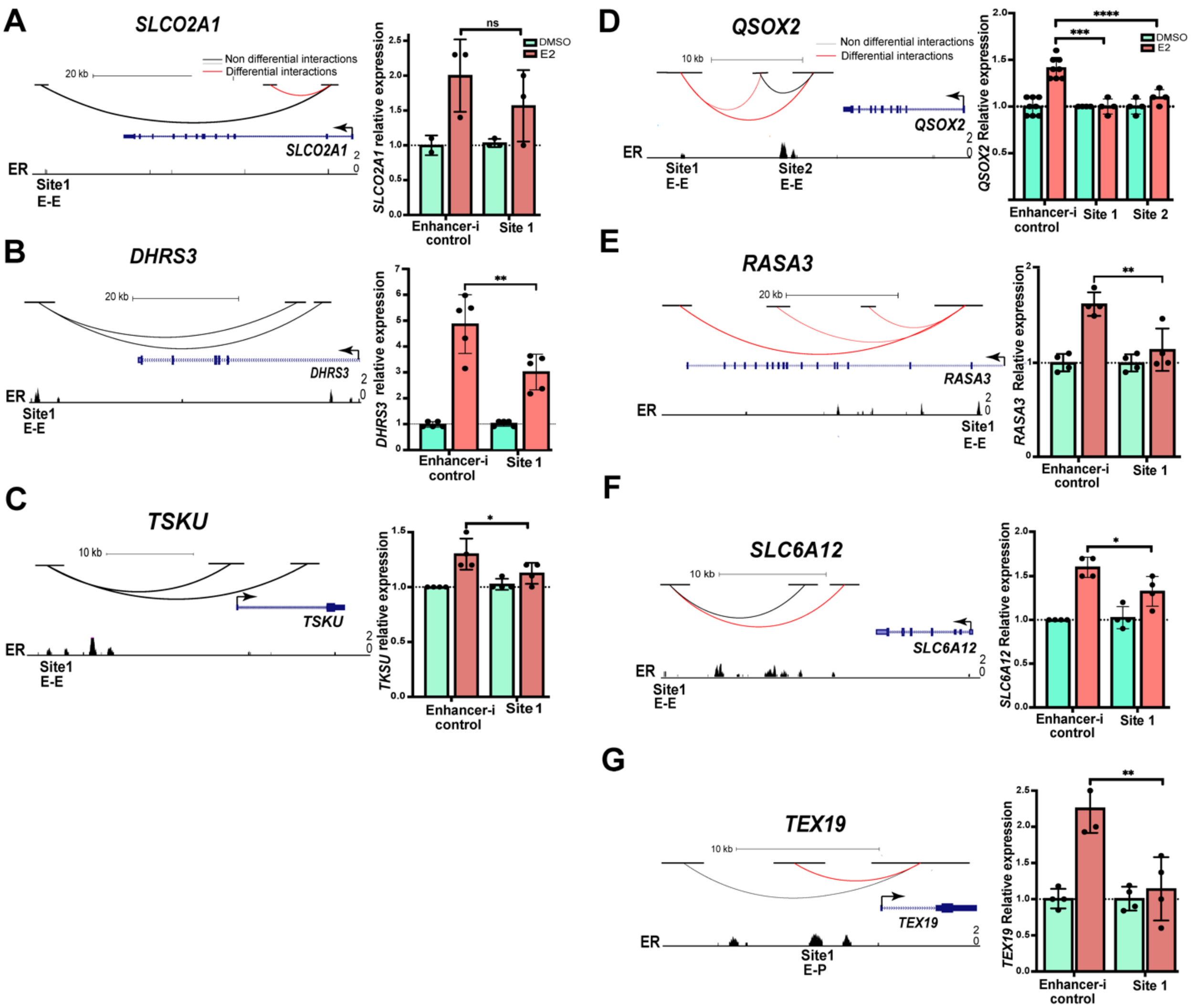


**Supplementary Figure 4 (related to Figure 4). ERBS that serve as chromatin loop anchors are functionally important. (**A-G) (left) Genome browser tracks of genes associated with ERBS with non-differential ((A) *SLCO2A1,*(B) *DHRS3,* (C) *TSKU*) and differential ((D) *QSOX2,* (E) *RASA3,* (F) *SLC6A12,* (G) *TEX19*) interactions are shown; red and black loops represent differential or non-differential interactions, respectively, and types of interactions are indicated by E-E or E-P under the browser track. ER ChIP-seq peaks are displayed at the bottom. (right) Relative expression of (A) *SLCO2A1,* (B) *DHRS3,* (C) *TSKU*, (D) *QSOX2,* (E) *RASA3,* (F) *SLC6A12,* and (G) *TEX19* are shown when an ERBS is targeted with Enhancer-i and cells are treated with DMSO (green) or E2 (orange). Error bars represent the SEM; ****p < 0.00001, ***p < 0.001, **p < 0.01 and *p < 0.05, unpaired t-test, while ns depict statistical insignificance.

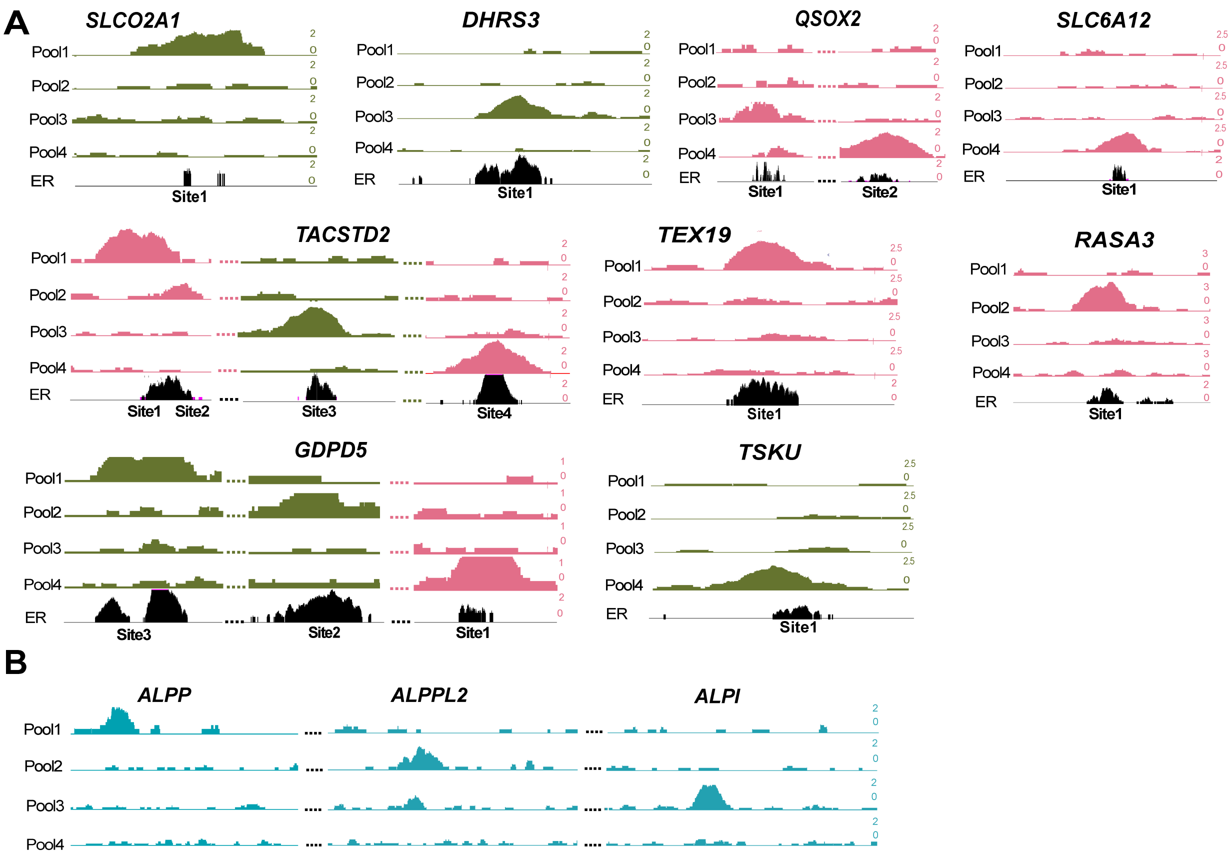


**Supplementary Figure 5 (related to Figure 4 and 5). Enhancer-i successfully targets promoters and enhancers.** A) ChIP-seq signal of Enhancer-i across pools of guide RNAs at targeted ERBS associated with differential (pink) or non-differential (green) interactions is shown along with ER ChIP-seq signal (black). B) ChIP-seq signal is shown for Enhancer-i across pools of guide RNAs at targeted promoters of genes that are part of an E2 upregulated gene cluster (*ALPP-APLPPL2-ALPI*).

### Table S1. Total targeted ERBS

| Site | Type of interaction | | Functional | % reduction | Study |
|---|---|---|---|---|---|
| TEX19 - Site2 | Promoter - Enhancer | Differential | Functional | 100 | This study |
| GDPD5 - Site1 | Promoter - Enhancer | Differential | Functional | 100 | This study |
| GDPD5 - Site2 | Enhancer-Enhancer | Non-differential | Functional | 100 | This study |
| GDPD5 - Site3 | Enhancer-Enhancer | Non-differential | Functional | 63 | This study |
| MMP17 - Site3 | Promoter - Enhancer | Differential | Non-functional | 12.9 | Carleton et al. |
| CISH - Site1 | Promoter - Enhancer | Differential | Functional | 44.8 | Carleton et al. |
| CISH - Site2 | Promoter - Enhancer | Differential | Functional | 42.2 | Carleton et al. |
| CISH - Site3 | Promoter - Enhancer | Non-differential | Non-functional | 3.9 | Carleton et al. |
| FHL2 - Site1 | Promoter - Enhancer | Non-differential | Functional | 64.5 | Carleton et al. |
| FHL2 - Site2 | Promoter - Enhancer | Non-differential | Functional | 42.5 | Carleton et al. |
| FHL2 - Site3 | Promoter - Enhancer | Non-differential | Non-functional | 14.3 | Carleton et al. |
| G0S2 - Site1 | Promoter - Enhancer | Non-differential | Functional | 93.3 | Carleton et al. |
| MMP17 - Site1/2 | Promoter - Enhancer | Non-differential | Functional | 33.7 | Carleton et al. |
| RASA3 - Site1 | Enhancer-Enhancer | Differential | Functional | 77.5 | This study |
| QSOX2 - Site1 | Enhancer-Enhancer | Differential | Functional | 100 | This study |
| QSOX2 - Site2 | Enhancer-Enhancer | Differential | Functional | 75.8 | This study |
| SLC6A12 - Site2 | Enhancer-Enhancer | Differential | Functional | 50 | This study |
| TACSTD2 - Site1 | Enhancer-Enhancer | Differential | Non-functional | 6.2 | This study |
| TACSTD2 - Site2 | Enhancer-Enhancer | Differential | Functional | 51.9 | This study |
| TACSTD2 - Site3 | Enhancer-Enhancer | Non-differential | Non-functional | 0 | This study |
| TACSTD2 - Site4 | Enhancer-Enhancer | Differential | Functional | 88.3 | This study |
| DHRS3 - Site1 | Enhancer-Enhancer | Non-differential | Functional | 48 | This study |
| SLCO2A1 - Site1 | Enhancer-Enhancer | Non-differential | Non-functional | 46.7 | This study |
| TKSU - Site1 | Enhancer-Enhancer | Non-differential | Functional | 66.7 | This study |

### Table S2. gRNA sequences

| gRNA name | gRNAs sequence | PAM |
|---|---|---|
| GDPD5 Enhancer gRNA 1 | AGAGAGCAGGAAGCAAGTTG | GGG |
| GDPD5 Enhancer gRNA 2 | CCAGTGTCCCCATATGACAT | AGG |
| GDPD5 Enhancer gRNA 3 | GTTCTCCCTGTCATCCCAAG | AGG |
| GDPD5 Enhancer gRNA 4 | GTAAACTGTACAGTGTGCAG | TGG |
| TEX19 Enhancer gRNA 1 | TCTTACCTGTGTCCTCGGGA | TGG |
| TEX19 Enhancer gRNA 2 | GAATGCCTCGCGCAGATCCG | GGG |
| TEX19 Enhancer gRNA 3 | GGTGCTCAAAGAGCGTGACG | TGG |
| TEX19 Enhancer gRNA 4 | TGCACCCCCGAGGTAGCTGG | GGG |
| RASA3 Enhancer gRNA 1 | GAATAAGAAGGGCGATTCCA | GGG |
| RASA3 Enhancer gRNA 2 | GCTCCCTGGCTATGGCATCG | TGG |
| RASA3 Enhancer gRNA 3 | GCCAGGGCGCTCAGCACGAG | GGG |
| RASA3 Enhancer gRNA 4 | GCTGCCATCCTAGCGCACAC | TGG |
| QSOX2 Enhancer 1 gRNA1 | TTAGGTAACCAGAGGCTGAT | GGG |
| QSOX2 Enhancer 1 gRNA2 | CCCCAAGAGCTCATGTGTCG | GGG |
| QSOX2 Enhancer 1 gRNA3 | TATTCCCGGGACCGGACTGG | AGG |
| QSOX2 Enhancer 1 gRNA4 | AGGTGCCAGGCCTTGGTCCG | AGG |
| QSOX2 Enhancer 2 gRNA1 | CAGTGCCAGCGTATATGCGA | GGG |
| QSOX2 Enhancer 2 gRNA2 | AGGCCAGGCCGACACCTCAA | TGG |
| QSOX2 Enhancer 2 gRNA3 | CTGGTGAGGTCGACGCGCTG | TGG |
| QSOX2 Enhancer 2 gRNA4 | CGCCTCTCCTGTTATCCGAG | GGG |
| SLC6A12 Enhancer 1 gRNA1 | GGGAGAGGTGCCCATTCCCG | TGG |
| SLC6A12 Enhancer 1 gRNA2 | AATCTCATGAGCTTTAGTGG | TGG |
| SLC6A12 Enhancer 1 gRNA3 | TGGTTGGAGAGTGTTCCTCG | TGG |
| SLC6A12 Enhancer 1 gRNA4 | TCACTCCAGCCGAGTCACAC | TGG |
| DHRS3 Enhancer gRNA1 | ACCCCATCTAATACGCACCA | TGG |
| DHRS3 Enhancer gRNA2 | GGGGATGTGGCCGTGAACTT | GGG |
| DHRS3 Enhancer gRNA3 | AGTTCTTAAAGGCCAACTGT | AGG |
| DHRS3 Enhancer gRNA4 | TTTTGTCCTCAAGGGGTAGC | AGG |
| TSKU Enhancer gRNA1 | CCAGCACAGGTGAACTGAAT | GGG |
| TSKU Enhancer gRNA2 | GTGCTACTCAGCACGAGCAG | AGG |
| TSKU Enhancer gRNA3 | AAAGAGGAGGGCCCGATCGA | GGG |
| TSKU Enhancer gRNA4 | GCAAAGCCAGCATTTGCTGA | GGG |
| SLCO2A1 Enhancer gRNA1 | ACACTCAGACTAGGCAGCAT | GGG |
| SLCO2A1 Enhancer gRNA2 | CTACCTGCTTCACTAGGAGT | CGG |
| SLCO2A1 Enhancer gRNA3 | CAGAAACCCCCGTACTTGAC | CGG |
| SLCO2A1 Enhancer gRNA4 | CACTGCACTTGGGTGTCCAA | GGG |
| ALPP Promoter gRNA1 | AGGGCCCCAGCATGTCTGGA | GGG |
| ALPP Promoter gRNA2 | TGTTGTGTAGGGGCAGCTCG | GGG |
| ALPP Promoter gRNA3 | GCAGGGTCAAGGTGGCAACG | AGG |
| ALPP Promoter gRNA4 | TCAATACCGCACCCTTCCCT | GGG |
| ALPPL2 Promoter gRNA1 | ATGGTGGATGAACGAGTGAC | AGG |
| ALPPL2 Promoter gRNA2 | AGCGGCGAGGCGGAAATCCC | AGG |
| ALPPL2 Promoter gRNA3 | TGGCTCACTCACCGTCACCC | AGG |
| ALPPL2 Promoter gRNA4 | CGACTGCTTCCAGACATGCA | GGG |
| ALPI Promoter gRNA1 | GACATTACGAGCCCTAACCC | GGG |

| | | |
|---|---|---|
| ALPI Promoter gRNA2 | GGCACCCATATACACCAAGT | GGG |
| ALPI Promoter gRNA3 | CGGGGTCAAGATGGACACCA | GGG |
| ALPI Promoter gRNA4 | TCCTCCCCTGATTTAAACCC | AGG |

### Table S3. qPCR primers

| Gene name | Forward sequence | Reverse sequence |
|---|---|---|
| CTCF | ACCTGTTCCTGTGACTGTACC | ATGGGTTCACTTTCCGCAAGG |
| GDPD5 | CCCATGATGATACCACACCGT | CCAGCGATGTATGCGAAGG |
| TEX19 | TCTACGCCTCCTGGATGTATC | CAGACCTGCATCTTCCAACCC |
| RASA3 | AGATCAAGATCGGTGAAGCC | CTCCGTAAAACGGGCAGAGT |
| QSOX2 | CTCGTGCAGTTCTACTCGTCG | AGCTCTCGGTCAGGTCCTTT |
| SLC6A12 | GGCGGACCGTTTCTATGACA | AGGAGGGTGATGACGACGAA |
| TACSTD2 | CGGCAGAACACGTCTCAGAAG | CCTTGATGTCCCTCTCGA AGTAG |
| DHRS3 | TCTGTGATGTGGGCAACCG | ATGGTGATGTCACCCACCTTC |
| TSKU | TCCAGCCTAGCCAGTTTCTC | TGTTGTGCTGCTTGTGACA |
| SLCO2A1 | GATGTCCACTGTCACCAAGC | GAATGACCTCCTTCGTTGGC |
| ALPP | CCCGCTTTAACCAGTGCAAC | GAGCTGCGTAGCGATGTCC |
| ALPPL2 | GGCATCATCCCAGTTGAGGAG | GCACATGCTTGTCTACACTGTAT |
| ALPI | TACACGTCCATCCTGTACGG | CTCGCTCTCATTCACGTCTGG |

### Table S4. Datasets used in the Boruta analyses

| Genomic Feature | Reference | Accession |
| --- | --- | --- |
| ATAC | (Vahrenkamp et al. 2018) | GSE109890 |
| CREB1 | (ENCODE 2012) | GSE32465 |
| CTCF | (ENCODE 2012) | GSE32465 |
| DNase | (ENCODE 2012) | GSE32970 |
| EGR1 | (ENCODE 2012) | GSE32465 |
| ER | (ENCODE 2012; Rodriguez et al. 2020) | GSE32465; GSE129803 |
| ETV4 | (ENCODE 2012) | GSE32465 |
| FOXA1 | (ENCODE 2012) | GSE32465 |
| FOXM1 | (ENCODE 2012) | GSE32465 |
| GR | (Vahrenkamp et al. 2018) | GSE109893 |
| H3K27ac | (Vahrenkamp et al. 2018) | GSE109893 |
| H3K27me3 | (Ginley-Hidinger et al. 2024) | GSE227241 |
| H3K36me3 | (Ginley-Hidinger et al. 2024) | GSE227241 |
| H3K4me1 | (Ginley-Hidinger et al. 2024) | GSE227241 |
| H3K4me3 | (Ginley-Hidinger et al. 2024) | GSE227241 |
| H3K9me3 | (Carleton et al. 2017) | GSE99906 |
| JUN | (Ginley-Hidinger et al. 2024) | GSE227241 |
| LSD1 | (Ginley-Hidinger et al. 2024) | GSE227241 |
| MAX | (Ginley-Hidinger et al. 2024) | GSE227241 |
| MYC | (Ginley-Hidinger et al. 2024) | GSE227241 |
| NFIC | (ENCODE 2012) | GSE32465 |
| NRSF | (ENCODE 2012) | GSE32465 |
| P300 | (ENCODE 2012) | GSE32465 |
| PRO-seq | (Ginley-Hidinger et al. 2024) | GSE227243 |
| RAD21 | (ENCODE 2012) | GSE32465 |
| RARA | (Ginley-Hidinger et al. 2024) | GSE227241 |
| RNAPII | (ENCODE 2012; Carleton et al. 2017) | GSE32465; GSE99906 |
| SIN3A | (Ginley-Hidinger et al. 2024) | GSE227241 |
| TAF1 | (Ginley-Hidinger et al. 2024) | GSE227241 |
| TCF12 | (ENCODE 2012) | GSE32465 |
| TEAD4 | (ENCODE 2012) | GSE32465 |
| USF1 | (ENCODE 2012) | GSE32465 |
| YY1 | (ENCODE 2012) | GSE32465 |
| ZBTB7A | (ENCODE 2012) | GSE32465 |
